# Tibial nerve stiffness is related to maximum angle of ankle dorsiflexion

**DOI:** 10.1101/2025.07.20.665743

**Authors:** Hiyu Mukai, Jun Umehara, Junya Saeki, Ko Yanase, Zimin Wang, Hiroshige Tateuchi, Noriaki Ichihashi

## Abstract

The maximum angle of ankle dorsiflexion is affected by the triceps surae muscle stiffness and stretch tolerance, which may be strongly reflected by the tibial nerve stiffness. However, no study has evaluated the triceps surae muscle and tibial nerve stiffness simultaneously or clarified their association with the maximum angle. The purpose of this study was to investigate the association between the maximum angle and both the stiffness. Forty-one healthy adults participated. The shear wave velocities of the triceps surae muscles and tibial nerve were measured at 5°, 15°, and 25° ankle dorsiflexion. Multiple linear regression analysis was performed using the forced entry method, specifying the shear wave velocities of the four tissues as the independent variables and the maximum angle as the dependent variable. This analysis was performed at each angle where the shear wave velocity was measured. Multiple linear regression analysis was also performed using the stepwise method, specifying the shear wave velocities of the tibial nerve at the three angles as the independent variables and the maximum angle as the dependent variable. Using the forced entry method, the shear wave velocity of the tibial nerve at each angle was significantly negatively associated with the maximum angle, whereas those of the muscles were not. Using the stepwise method, only the shear wave velocity of the tibial nerve at 25° was significantly negatively associated with the maximum angle. These results suggest that the tibial nerve stiffness in a greatly lengthened position determines the maximum angle of ankle dorsiflexion.

## 1. Introduction

The maximum angle of ankle dorsiflexion is an index of joint flexibility that is frequently used in clinical assessments. It is associated with impaired balance and walking ability in elderly individuals (Chiara Mecagni et al., 2000) and is a risk factor for patellar tendinopathy (Backman and Danielson, 2011; Malliaras et al., 2006). Thus, it is important to investigate the factors that determine the maximum angle of ankle dorsiflexion.

Conventionally, the maximum joint angle is considered to be affected by muscle stiffness and stretch tolerance (Magnusson et al., 1997). Recently, nerve stiffness was suggested to be associated with stretch tolerance (Andrade et al., 2018). Although some previous studies have reported a correlation between muscle or nerve stiffness and the maximum angle of ankle dorsiflexion, few studies have measured muscle and nerve stiffness simultaneously and investigated their correlations with the maximum angle of ankle dorsiflexion. The medial and lateral gastrocnemius muscle stiffness of healthy young males at 14° of ankle dorsiflexion were shown to be negatively correlated with the maximum angle of ankle dorsiflexion (Miyamoto et al., 2018). However, the findings of previous studies regarding the association between the lateral gastrocnemius muscle stiffness assessed in a slightly lengthened position (i.e., at 0° of ankle dorsiflexion) and maximum angle of ankle dorsiflexion are inconsistent (Miyamoto et al., 2018; Reiner et al., 2024). Additionally, these studies did not evaluate nerve stiffness. Kawanishi et al. (2022) reported a negative correlation between the tibial nerve stiffness at 75% and 100% of the maximum angle of ankle dorsiflexion and maximum angle of ankle dorsiflexion itself; however, they did not evaluate muscle stiffness. These studies suggested that muscle and nerve stiffness may be associated with the maximum angle of ankle dorsiflexion. However, few studies have comprehensively investigated the association between both stiffness and the maximum angle of ankle dorsiflexion.

Hirata et al. (2020) reported that in healthy young males, the triceps surae muscle stiffness at 15° of ankle dorsiflexion was negatively correlated with the maximum angle of ankle dorsiflexion, whereas the sciatic nerve stiffness was not. However, the tibial nerve, which may affect the maximum angle of ankle dorsiflexion more strongly than the sciatic nerve, has not been investigated. The tibial nerve stiffness near the ankle joint is negatively correlated with the maximum angle of ankle dorsiflexion (Kawanishi et al., 2022) and is higher than the sciatic nerve stiffness(Andrade et al., 2022). Thus, it may reflect the stretch tolerance to ankle dorsiflexion more strongly than the sciatic nerve stiffness. However, no study has investigated the association between the triceps surae muscle and tibial nerve stiffness and maximum angle of ankle dorsiflexion, and it is unclear which stiffness is associated most strongly. In addition, stiffness at angles of ankle dorsiflexion larger than 15° may be more negatively associated with the maximum angle of ankle dorsiflexion.

The purpose of this study was to evaluate both triceps surae muscle and tibial nerve stiffness and to clarify their association with the maximum angle of ankle dorsiflexion using shear wave elastography. Negative correlations were found between i) the triceps surae muscle stiffness and maximum angle of ankle dorsiflexion(Hirata et al., 2020; Miyamoto et al., 2018) and ii) the tibial nerve stiffness and maximum angle of ankle dorsiflexion (Kawanishi et al., 2022). Based on these correlations, we hypothesized that these muscles and nerve stiffness were associated with the maximum angle of ankle dorsiflexion and that the association was stronger when the muscles and nerve were greatly lengthened.

## 2. Methods

### 2.1. Participants

Forty-three healthy young adults who had no pain or reduced range of motions in their ankles on the nondominant side participated in this study (19 males and 24 females, age 24.3 ± 2.9 years; height 165.8 ± 7.7 cm; body mass 59.3 ± 8.2 kg). Thirty-nine of them were right-footed and four were left-footed when the dominant foot was defined as the one that kicked a ball. An a priori sample size estimation was performed using a linear multiple regression model in G*Power (version 3.1.9.7; Heinrich Heine University, Düsseldorf, Germany) using an assumed effect size of 0.35, an alpha level of 0.05, and a statistical power of 0.80. The analysis indicated that a minimum of 40 participants were required. Forty-three participants were recruited to account for potentially missing data. The study protocols were explained to all the participants and informed consent was obtained from each participant. This study conformed to the Declaration of Helsinki and was approved by the Ethics Committee (approval number: C1652-2).

### 2.2. Experimental procedure

This cross-sectional study investigated the association between different shear wave velocities and the maximum angle of ankle dorsiflexion. The shear wave velocities of the medial and lateral gastrocnemius muscles, soleus muscle, and tibial nerve were measured. The participants were placed in a prone position on the seat of a dynamometer (BIODEX System 4, BIODEX, NY, USA) to measure the shear wave velocities and maximum angle. They were set at 0° of hip flexion/extension, abduction/adduction, and internal/external rotation and 0° of knee flexion/extension, with their feet attached to a plate and their feet and pelvis fixed using straps. The participants were asked to relax during the measurements. To minimize the stretching effect, first, the shear wave velocities were measured at 5°, 15°, and 25° of ankle dorsiflexion, in order, and then the maximum angle of ankle dorsiflexion was measured.

### 2.3. Measurement of shear wave velocity

The shear wave elastography mode in an ultrasound system (Aixplorer v12.2, SuperSonic Imagine, Aix-en-Provence, France) with a linear probe (2–10 MHz, SuperLinear SL10-2) was used to measure the shear wave velocity in each tissue. Each measurement was performed in a musculoskeletal preset (muscle mode) (penetration mode, frequency: 1.7 Hz, smoothing level: five, persistence high, opacity: 100%). Shear wave velocity (V) is used as an index of muscle and nerve stiffness and is directly related to the shear modulus (G) as follows:

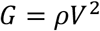

where ρ is the estimated density of tissues (1.0 g/cm^3^) (Nordez et al., 2008). Muscle shear modulus is strongly correlated with Young’s modulus, as assessed by traditional material testing (Eby et al., 2013). However, the density of the tibial nerve could not be estimated as 1.0 g/cm^3^, and we adopted the shear wave velocity in this study. We interpreted that a higher shear wave velocity indicates a higher stiffness of tissues. The shear wave velocities were measured at the following locations: the medial and lateral gastrocnemius muscles at 30% of the lower leg length (Akagi and Takahashi, 2013; Nakamura et al., 2014), soleus muscle at 50% of the lower leg length (Kubo et al., 2017), and tibial nerve at a point near the medial malleolus, as determined by preliminary experiments.

### 2.4. Image analysis

A region of interest (ROI) with a length of 1 cm and width of 2 cm was set. The Q-box trace function was then used to surround the maximal area in the ROI, excluding the fascia, epineurium, and artifacts, and the average shear wave velocity within the area was calculated. For each joint angle, two ultrasound images of the triceps surae muscles and three ultrasound images of the tibial nerve were acquired, and their mean values were used for statistical analysis. The analyzed ultrasound images are shown in Fig. 1.

**Fig. 1:**
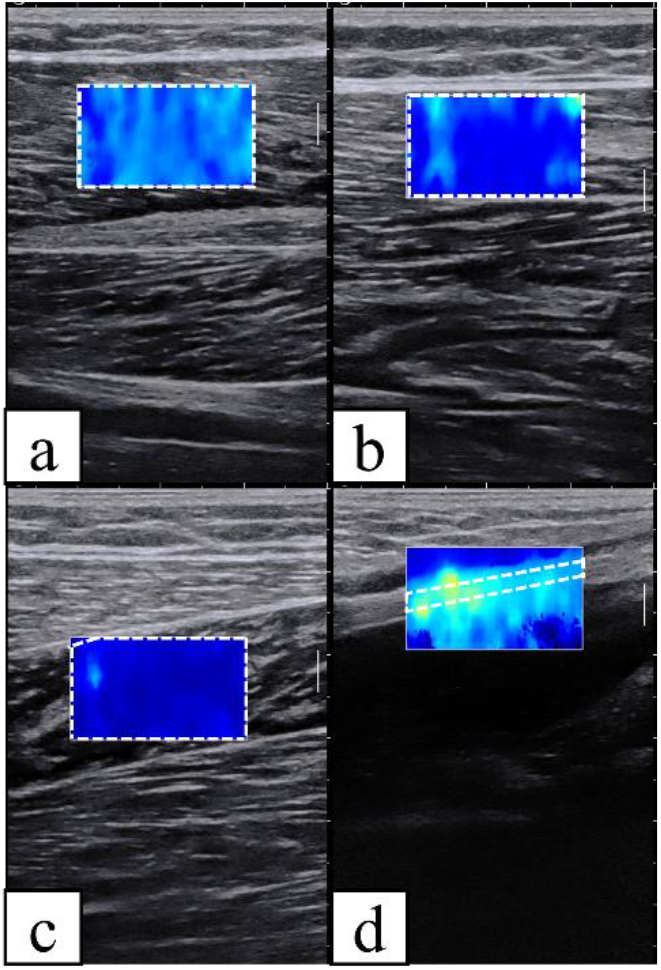
The analyzed ultrasound images. a: medial gastrocnemius muscle, b: lateral gastrocnemius muscle, c: soleus muscle, d: tibial nerve. The areas surrounded by white dashed lines are analyzed.

### 2.5. Measurement of maximum angle of ankle dorsiflexion

The ankle of a participant was passively dorsiflexed at 5°/s velocity from 30° of ankle plantar flexion to the maximum angle at which the participant experienced discomfort without pain (Nakamura et al., 2017). The participants were asked to press a button to stop the footplate upon reaching the maximum angle of ankle dorsiflexion. The maximum angle was defined in 1-degree increments. Before the measurement of the maximum angle of ankle dorsiflexion, two sessions were conducted to familiarize the participants with the procedure(Hirata et al., 2015; Konrad et al., 2015; Konrad and Tilp, 2014). The maximum angle of ankle dorsiflexion was measured three times, and the mean value was used for statistical analysis.

### 2.6. Statistical analysis

One participant who felt pain while measuring the shear wave velocities at 25° of ankle dorsiflexion and another participant whose maximum angle of ankle dorsiflexion exceeded the upper limit of the dynamometer were excluded. We performed statistical analysis of the data obtained from 41 participants. SPSS Statistics 22 (IBM, Armonk, NY, USA) was used for the statistical analysis. To confirm whether each tissue lengthened with increasing ankle dorsiflexion, a multiple comparison test with Bonferroni correction was performed to compare the shear wave velocities at 5°, 15°, and 25° of ankle dorsiflexion. To investigate the association between the shear wave velocities of each tissue and maximum angle of ankle dorsiflexion, a multiple linear regression analysis using the forced entry method was performed, specifying the shear wave velocities of the four tissues as the independent variables and the maximum angle of ankle dorsiflexion as a dependent variable. This multiple linear regression model was built at each angle where the shear wave velocity was measured because previous studies have reported inconsistent findings regarding the association between the shear wave velocity of a muscle in a slightly lengthened position and maximum angle(Miyamoto et al., 2018; Reiner et al., 2024). Additionally, to investigate the angle at which the shear wave velocity measured was most associated with the maximum angle, a multiple linear regression analysis using the stepwise method was performed, specifying the shear wave velocities that were significantly associated with the maximum angle of ankle dorsiflexion in the forced entry method as the independent variables and the maximum angle of ankle dorsiflexion as a dependent variable. Statistical significance was set as *p* < 0.05.

## 3. Results

### 3.1. Characteristics and shear wave velocity of participants

The characteristics of the 41 participants included in the analysis were shown as follows; 19 males and 22 females, 37 right-footed and 4 left-footed, age—24.3 ± 2.9 years; height—166.2 ± 7.6 cm; body mass—59.8 ± 7.6 kg. The mean ± SD of their maximum angle of ankle dorsiflexion was 32 ± 7°. For the medial and lateral gastrocnemius muscles, the shear wave velocities at 15° of ankle dorsiflexion were significantly higher than those at 5° of ankle dorsiflexion, and the shear wave velocities at 25° of ankle dorsiflexion were significantly higher than those at 5° and 15° of ankle dorsiflexion (*p* < 0.001 for all the angles). For the soleus muscle, no significant difference was found in the shear wave velocities at 5° and 15° of ankle dorsiflexion (*p* = 0.557), whereas the shear wave velocity at 25° of ankle dorsiflexion was significantly higher than those at 5° and 15° of ankle dorsiflexion (both *p* < 0.001). For the tibial nerve, the shear wave velocity at 15° of ankle dorsiflexion was significantly higher than that at 5° of ankle dorsiflexion (*p* = 0.001), and the shear wave velocity at 25° of ankle dorsiflexion was significantly higher than those at 5° (*p* < 0.001) and 15° (*p* = 0.002) of ankle dorsiflexion. The shear wave velocity of each tissue at each angle is listed in Table 1.

**Table 1:** Shear wave velocity of each tissue at each angle.

|  | Shear wave velocity (m/s) |  |  |
| --- | --- | --- | --- |
|  | 5° of ankle dorsiflexion | 15° of ankle dorsiflexion | 25° of ankle dorsiflexion |
| Medial gastrocnemius muscle | 4.59 ± 0.61 | 6.45 ± 0.78* | 8.32 ± 0.82*§ |
| Lateral gastrocnemius muscle | 4.05 ± 0.66 | 5.25 ± 0.67* | 6.95 ± 0.88*§ |
| Soleus muscle | 3.31 ± 0.90 | 3.46 ± 0.75 | 4.00 ± 0.93*§ |
| Tibial nerve | 6.74 ± 1.31 | 7.24 ± 1.40* | 7.78 ± 1.23*§ |

Shear wave velocities are described as mean ± SD. \*: significantly higher than 5° of ankle dorsiflexion ( $p < 0.05$ ), §: significantly higher than 15° of ankle dorsiflexion ( $p < 0.05$ ).

### 3.2. Multiple linear regression analysis of shear wave velocity and maximum angle of ankle dorsiflexion

The results of the multiple linear regression analysis using the forced entry method are presented in Table 2. The shear wave velocities of the tibial nerve at 5° (adjusted R^2^ = 0.045), 15° (adjusted R^2^ = 0.177), and 25° (adjusted R^2^ = 0.138) of ankle dorsiflexion were significantly and negatively associated with the maximum angle of ankle dorsiflexion. However, the shear wave velocities of the triceps surae muscles at 5°, 15°, and 25° of ankle dorsiflexion were not significantly associated with the maximum angle of ankle dorsiflexion. Multiple linear regression analysis was also conducted using the stepwise method, specifying the shear wave velocities of the tibial nerve at 5°, 15°, and 25° of ankle dorsiflexion as the independent variables. Its results showed that only the shear wave velocity of the tibial nerve at 25° of ankle dorsiflexion was significantly and negatively associated with the maximum angle of ankle dorsiflexion (unstandardized regression coefficient = −2.618, standardized regression coefficient = −0.449, 95% confidence interval = from −4.305 to −0.931, *p* = 0.003, adjusted R^2^ = 0.181).

**Table 2:** The results of multiple regression analysis.

| 5° of ankle dorsiflexion |  |  |  |  |
| --- | --- | --- | --- | --- |
|  | B | β | 95%CI | *p* value |
| Medial gastrocnemius muscle | 0.955 | 0.081 | -3.362 to 5.272 | 0.656 |
| Lateral gastrocnemius muscle | -0.939 | -0.086 | -5.117 to 3.298 | 0.656 |
| Soleus muscle | 0.054 | 0.007 | -2.715 to 2.822 | 0.969 |
| Tibial nerve | -1.977 | -0.361 | -3.759 to -0.195 | 0.031* |
| 15° of ankle dorsiflexion |  |  |  |  |
|  | B | β | 95%CI | *p* value |
| Medial gastrocnemius muscle | -3.174 | -0.345 | -6.908 to 0.559 | 0.093 |
| Lateral gastrocnemius muscle | 3.728 | 0.347 | -1.194 to 8.650 | 0.133 |
| Soleus muscle | 1.355 | 0.141 | -1.690 to 4.399 | 0.373 |
| Tibial nerve | -2.954 | -0.575 | -4.719 to -1.188 | 0.002* |
| 25° of ankle dorsiflexion |  |  |  |  |
|  | B | β | 95%CI | *p* value |
| Medial gastrocnemius muscle | -0.808 | -0.092 | -3.686 to 2.071 | 0.573 |
| Lateral gastrocnemius muscle | 1.101 | 0.134 | -1.671 to 3.874 | 0.426 |
| Soleus muscle | 0.582 | 0.075 | -1.844 to 3.008 | 0.630 |
| Tibial nerve | -2.880 | -0.494 | -4.702 to -1.059 | 0.003* |

\*: $p < 0.05$ , B: unstandardized regression coefficient, β: standardized regression coefficient, CI: confidence interval.

## 4. Discussion

In this study, we investigated the association between the shear wave velocities of the triceps surae muscles and tibial nerve and the maximum angle of ankle dorsiflexion. We found that the shear wave velocities of the tibial nerve at 5°, 15°, and 25° of ankle dorsiflexion were negatively associated with the maximum angle and that the shear wave velocity of the tibial nerve at 25° of ankle dorsiflexion was the most negatively associated. However, the shear wave velocities of the triceps surae muscles measured at these angles were not associated with the maximum angle. This is the first study to show that the tibial nerve stiffness is more strongly associated with the maximum angle of ankle dorsiflexion than the triceps surae muscle stiffness.

The significant association between the shear wave velocity of the tibial nerve and maximum angle of ankle dorsiflexion is aligned with our hypothesis and the findings of a previous study(Kawanishi et al., 2022). Andrade et al. (2018) suggested that nerve stiffness reflects stretch tolerance and that its potential mechanism is an intrinsic nerve such as the nervi nervorum. The nervi nervorum may be related to pain threshold (Bove and Light, 1995; Marshall, 1883) and is sensitive to stretching along the long axis of the nerve (Andrade et al., 2018). Individuals with high tibial nerve stiffness may have poor stretch tolerance to ankle dorsiflexion owing to a sensitive nervi nervorum. However, the mechanism underlying the effect of nerve stiffness on stretch tolerance remains unclear. Additionally, using the stepwise method, we found that only the shear wave velocity of the tibial nerve at 25° of ankle dorsiflexion was associated with the maximum angle of ankle dorsiflexion. The tibial nerve stiffness in a greatly lengthened position may be more strongly associated with the maximum angle of ankle dorsiflexion than that in a slightly lengthened position. The greatly lengthened nervi nervorum of the tibial nerve may reflect stretch tolerance to ankle dorsiflexion, although the underlying mechanism has not been clarified.

The nonsignificant association between the shear wave velocities of the triceps surae muscles and maximum angle of ankle dorsiflexion was not aligned with our hypothesis and previous studies (Hirata et al., 2020; Miyamoto et al., 2018). Conventionally, both muscle stiffness and stretch tolerance are considered to be associated with the maximum joint angle (Magnusson et al., 1997). Previous studies have reported a nonsignificant association between the maximum angle of ankle dorsiflexion and medial and lateral gastrocnemius muscle stiffness in a slightly lengthened position (Reiner et al., 2024) and a significant association between them in a greatly lengthened position (Hirata et al., 2020; Miyamoto et al., 2018). However, in this study, the shear wave velocities of the triceps surae muscles, even in greatly lengthened positions, were not associated with the maximum angle of ankle dorsiflexion. This study suggested that stretch tolerance affects the maximum joint angle more strongly than muscle stiffness. Nakamura et al. (2021) reported that an increase in the maximum angle of ankle dorsiflexion did not correlate with a decrease in the medial gastrocnemius muscle stiffness after static stretching intervention. Although this was a cross-sectional study and not an intervention study, muscle stiffness may be less strongly associated with the maximum joint angle than stretch tolerance.

This study had some limitations. First, the participants were healthy young adults. Hirata et al. (2020) reported that tissue stiffness correlation with the maximum joint angle differed between young and elderly people. Additionally, because the gastrocnemius muscle stiffness of stroke patients is higher than that of healthy individuals (Belghith et al., 2024; Le Sant et al., 2019), in such patients, muscle stiffness may be associated with the maximum joint angle. Second, the shear wave velocities of the triceps surae muscles and tibial nerve were measured at a single point. Previous studies reported regional differences in the medial and lateral gastrocnemius muscles (Le Sant et al., 2017; Zhou et al., 2019) and tibial nerve stiffness (Andrade et al., 2022). A region where the tissue stiffness is more strongly associated with the maximum joint angle may be found by measuring the shear wave velocities at other points. Third, we measured the shear wave velocities and maximum angles with only 0° of knee flexion. Previous studies have reported that when the ankle moves from plantarflexion to dorsiflexion, the change in the triceps surae muscle stiffness differs for 0° and 90° of knee flexion (Ateş et al., 2018; Le Sant et al., 2017). The maximum angle of ankle dorsiflexion with different angle of knee flexion may be associated with tissue stiffness other than that of the tibial nerve.

Consequently, the tibial nerve stiffness was negatively associated with the maximum angle of ankle dorsiflexion, whereas the triceps surae muscle stiffness was not. Because nerve stiffness is considered to reflect stretch tolerance (Andrade et al., 2018), stretch tolerance may be more strongly associated with the maximum angle of ankle dorsiflexion than muscle stiffness. Additionally, the tibial nerve stiffness in a greatly lengthened position was more strongly associated with the maximum angle of ankle dorsiflexion than in a slightly lengthened position. Nerve stiffness in a greatly lengthened position may more strongly reflect stretch tolerance.

## Declaration of Ethics

All relevant ethical guidelines have been followed, all necessary ethics committee approvals have been obtained, all necessary participant consent has been obtained, and the appropriate institutional forms have been archived.

## Acknowledgement

We are grateful to Editage (www.editage.jp) for assistance with English language editing.

